# Parasites as niche modifiers for the microbiome: A field test with multiple parasites

**DOI:** 10.1101/2020.03.31.018713

**Authors:** Kayleigh R. O’Keeffe, Fletcher W. Halliday, Corbin D. Jones, Ignazio Carbone, Charles E. Mitchell

## Abstract

Parasites can affect and be affected by the host’s microbiome, with consequences for host susceptibility, parasite transmission, and host and parasite fitness. Yet, there are two aspects of the relationship between parasite infection and the host microbiome that remain little understood: the nature of the relationship under field conditions, and how the relationship varies among parasite species. To overcome these limitations, we assayed the within-leaf fungal community in a grass population to investigate how diversity and composition of the fungal microbiome are associated with natural infection by fungal parasites with different feeding strategies. We hypothesized that parasites that more strongly modify niches available within a host will thereby alter the microbial taxa that can colonize the community and be associated with greater changes in microbiome diversity and composition. A parasite that creates necrotic tissue to extract resources (necrotrophs) may act as a particularly strong niche modifier whereas one that does not (biotrophs) may not. Barcoded amplicon sequencing of the fungal ITS region revealed that the microbiome of leaf segments that were symptomatic of necrotrophs had lower fungal diversity and distinct composition compared to segments that were asymptomatic or symptomatic of other parasites. There were no clear differences in fungal diversity or composition between leaf segments that were asymptomatic and segments that were symptomatic of other parasite feeding strategies. This supports the hypothesis that within-host niches link infection by parasites to the host’s microbiome. Together, these results highlight the importance of parasite traits in determining parasite impacts on the host’s microbiome.

## Introduction

The fungi, bacteria, and viruses that comprise an organism’s microbiome can interact with parasites in ways that have consequences for host susceptibility, disease severity, and transmission (Berg & Koskella, 2018). Yet, while investigations of the association between the microbiome and a single parasite species are becoming increasingly common (Aivelo & Norberg, 2018; Libertucci & Young, 2019), how such associations vary among parasite species is still poorly understood. Here, we evaluated potential mechanisms through which parasites alter the fungal microbiome using a trait-based analysis of multiple co-occurring fungal parasites in a grass host. Specifically, we test the hypothesis that parasites with a trait that more strongly modifies the environment within the host act as niche modifiers and more strongly impact microbiome diversity and composition.

Microbes may interact with parasites by competing for resources, releasing antimicrobial compounds, or eliciting a host immune response (Graham, 2008; Raaijmakers & Mazzola, 2012; Bashey, 2015). Such interactions can influence host health, making the host more susceptible to, tolerant of, or resistant to parasites (A. E. Arnold et al., 2003; Busby, Peay, & Newcombe, 2016; Hayes et al., 2010). The microbiome is also dynamic, and the introduction of a parasite can lead to a change in microbiome diversity and composition (Barman et al., 2008; Jani & Briggs, 2014). The link between parasites and host microbial diversity varies, with some studies showing no relationship (Li et al., 2012; Williams et al., 2017), some studies showing a negative relationship (Jani & Briggs, 2014; Leung et al., 2018; Mosca, Leclerc, & Hugot, 2016; Wu, Stanley, Rodgers, Swick, & Moore, 2014), and still others showing a positive relationship between parasite infection and within-host microbial diversity (Lee et al., 2014). One possible source of this variation among studies is variation among the parasite species studied. Yet, there is a lack of studies that examine variation among parasite species in their associations with host microbiota (but see Aivelo & Norberg, 2018), perhaps because we have lacked a robust framework for studying multiple parasites.

A trait-based approach might provide a framework for studying multiple parasite species (Zanne et al., 2019). As well as being evolutionarily diverse (Weinstein & Kuris, 2016), parasites vary in traits such as growth rate, generation time, and feeding strategy (Leggett, Cornwallis, Buckling, & West, 2017). Different parasite feeding strategies can stimulate different immune responses in a host and differently impact host performance (Glazebrook, 2005; Budischak, O’Neal, Jolles, & Ezenwa, 2018; Halliday, Umbanhowar, & Mitchell, 2018). Parasites infecting plants employ three typical feeding strategies. Parasites with a biotrophic feeding strategy keep host cells alive to extract resources from them. Parasites with a necrotrophic feeding strategy kill host cells to access their resources, creating necrotic tissue while they grow within their host. Finally, hemibiotrophic parasites infect as biotrophs, then switch to become necrotrophic. Given that parasites with different feeding strategies have different impacts on the host environment, parasites with different feeding strategies may also have different associations with the host microbiome.

Because feeding strategy can define how a parasite impacts the host environment, a trait-based approach grounded in parasite feeding strategy might help explain why some parasites have larger impacts on the within-host microbial community than other parasites. When an organism modifies its environment, it can change the number and types of niches available, and in turn, impact the species that can reside and colonize within the ecological community via the process of niche modification (Lewontin, 1983; Fukami, 2015). Niche modification is closely related to the concepts of niche construction (particularly with respect to evolutionary implications, Odling-Smee, Laland, & Feldman, 2003) and ecosystem engineering (Jones, Lawton, & Shachak, 1994), and has been documented in numerous communities of free-living organisms (Naiman et al., 2009; Fukami & Nakajima, 2011). Parasites may also act as niche modifiers within host individuals by impacting the host environment in such a way that the host is more or less suitable to new colonizers. Specifically, a parasite that produces necrotic tissue can create new niches within the host, allowing colonization by new taxa, and thus may lead to a particularly large change in microbiome richness and composition. A parasite that keeps host cells alive while extracting resources may have a more subtle impact on niches within the host, and thus may not change microbiome diversity or composition. Thus, the stability of the microbiome in the face of a parasite infection, which can impact disease severity and host health (Coyte, Schluter, & Foster, 2015), may be determined by parasite feeding strategy.

Few studies have investigated the associations between host-associated microbiota and parasites under field conditions (but see Jani & Briggs, 2014). This lack of studies may result from a limited number of suitable model systems for exploring these questions in the field. The long-lived nature of some hosts, limited ability to detect diseases observationally in live hosts in the field, difficulty of excising infected tissue from animals, and ethical concerns also limit the utility of many study systems for field research (Borer et al., 2011). As such, most investigations of parasite-microbiome relationships have been conducted under controlled settings (Leung et al., 2018). In a rare study in which the relationship between parasite infection and the host microbiome was studied in both the lab and the field, the direction and magnitude of the relationship differed between laboratory and field conditions (Leung et al., 2018). This finding underscores the importance of investigating parasite-microbiome associations in field settings.

Foliar fungal parasites are a valuable model system to investigate microbiome-parasite interactions in field settings. These parasites are often readily identifiable by external, macroscopic symptoms and morphology, which facilitates observational studies in the field. While bacteria can be numerically more abundant than fungi within plant hosts (Lundberg et al., 2012), fungi are functionally important symbionts of plants, with clear impacts on plant health (Christian, Whitaker, & Clay, 2015). Many studies have examined the relationship between the fungal microbiome and a single species of plant parasite (A. E. Arnold et al., 2003; Busby, Ridout, & Newcombe, 2016). In contrast, no plant microbiome studies to our knowledge have simultaneously considered multiple parasites in a common environment, either in the lab or in the field.

To fill this gap, we conducted a molecular survey of the within-leaf fungal community of the grass, tall fescue, in a field population infected by three fungal parasites. These three parasites differed in a key ecological trait: feeding strategy (biotrophic, hemibiotrophic, and necrotrophic). We hypothesized that necrotrophic parasites act as niche modifiers for the microbiome. More specifically, we hypothesized that necrotrophic parasites, by producing necrotic tissue, create new niches within the host, and thus have larger impacts on microbiome richness and composition than biotrophic parasites, which keep host cells alive while extracting resources. We further hypothesized that when hemibiotrophic parasites produce necrotic tissue, they, like necrotrophs, will act as niche modifiers, but to a lesser degree than necrotrophs because they only employ the necrotrophic feeding strategy in later stages of infection. This hypothesis predicts that the composition of the fungal community in leaves exhibiting symptoms of necrotrophic parasites will differ from the fungal communities in asymptomatic leaves and in leaves exhibiting symptoms of biotrophic parasites.

## Methods

### Study System

Leaf segments were collected in a grass-dominated field within the Blackwood Division of the Duke Forest Teaching and Research Laboratory in Orange County, North Carolina (35° 58’N, 79° 5’W). This field was chosen based on proximity to the lab (to expedite sample processing), and abundance of tall fescue (*Lolium arundinaceum*) and its foliar fungal parasites. This study focused on disease symptoms previously identified in another field of the Duke Forest Teaching and Research Laboratory (Halliday *et al*. 2017) as representing parasites with three different parasite feeding strategies (Table 1).

**Table 1.**
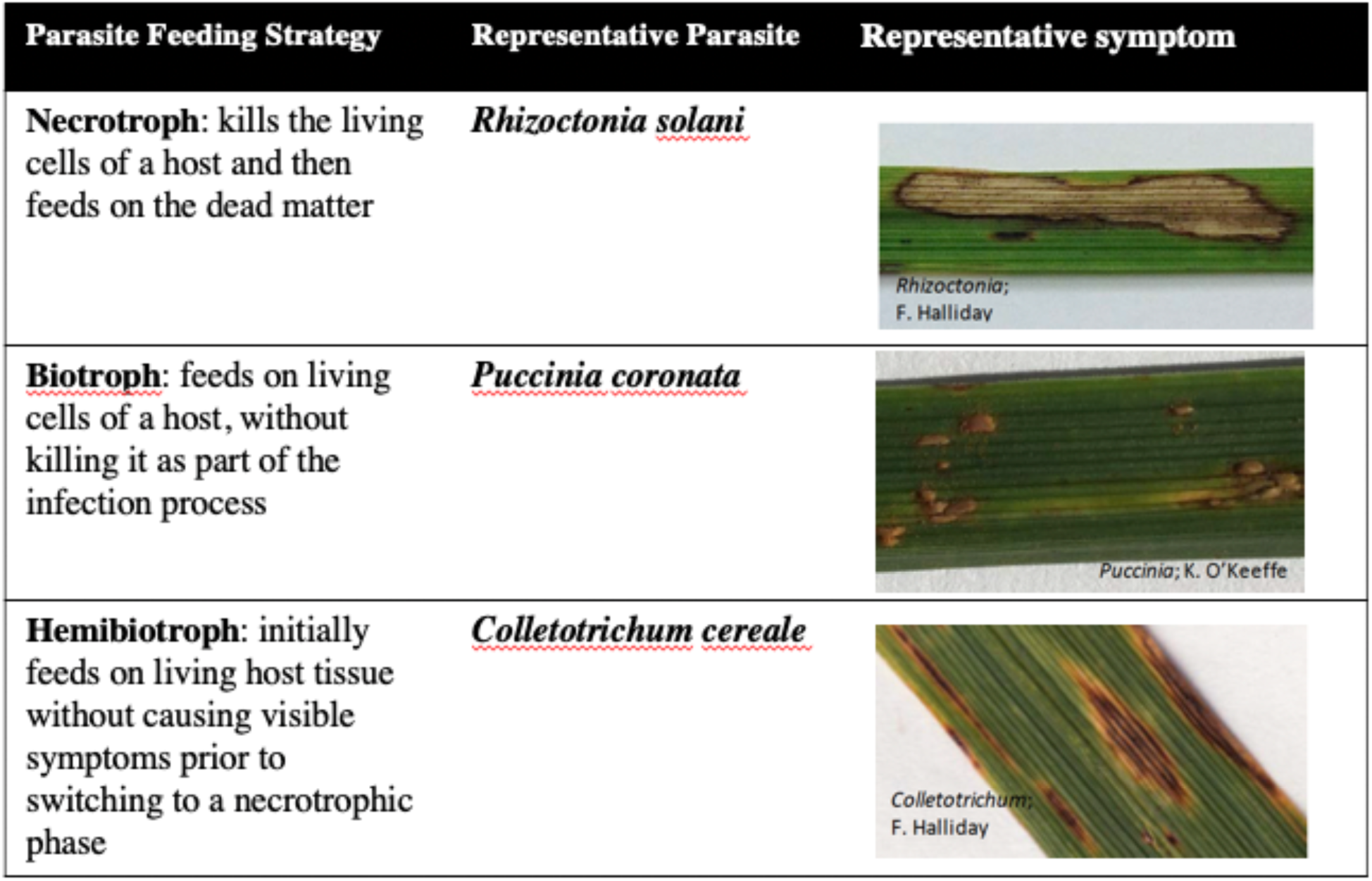
Biology and ecology of the focal parasite feeding strategies.

| Parasite Feeding Strategy | Representative Parasite | Representative symptom |
| --- | --- | --- |
| **Necrotroph:** kills the living cells of a host and then feeds on the dead matter | *Rhizoctonia solani* | *Rhizoctonia;*<br>F. Halliday |
| **Biotroph:** feeds on living cells of a host, without killing it as part of the infection process | *Puccinia coronata* | *Puccinia;* K. O'Keeffe |
| **Hemibiotroph:** initially feeds on living host tissue without causing visible symptoms prior to switching to a necrotrophic phase | *Colletotrichum cereale* | *Colletotrichum;*<br>F. Halliday |

### Leaf Segment Collection

Leaf segments were collected at a total of 36 points, spaced every 20 meters along 6 transects; the transects were 100m long, parallel, and spaced 20 meters apart (Figure S1). Leaf segments were collected over the course of two days in late September 2016 (September 26 and September 30), which is when parasites tend to peak in their abundance in this system (Halliday, Umbanhowar, & Mitchell, 2017).

At each point along each transect, we collected four whole leaves—one leaf with symptoms of a necrotrophic parasite (and no other parasites), one leaf with symptoms of a hemibiotrophic parasite (and no other symptoms), one leaf with symptoms of a biotrophic parasite (and no other parasites), and one asymptomatic leaf. While coinfection is common in this system (Halliday et al., 2017), we avoided collecting coinfected leaves. Only one leaf of each symptom was collected at each point, and therefore leaves with the same symptom were always collected at least 8 meters apart. This minimum distance was selected to minimize the effect of spatial autocorrelation on the structure and composition of the microbiome (A. E. Arnold, Henk, Eells, Lutzoni, & Vilgalys, 2007; Higgins, Arnold, Miadlikowska, Sarvate, & Lutzoni, 2007). At each sampling point, for each symptom, leaves were haphazardly collected by looking away, placing a finger on a plant, and then selecting the nearest tall fescue tiller. To standardize the relative age of sampled leaves, we always sampled the oldest fully expanded non-senescing leaf on the tiller. For each leaf, we estimated the percent of leaf area infected with each parasite (infection severity) by visually comparing leaves to reference images of leaves of known infection severity (James, 1971, e.g., Mitchell, Tilman, & Groth, 2002, 2003; Halliday, Heckman, Wilfahrt, & Mitchell, 2019).

At each of the 36 sampling points, we collected seven leaf segments: symptomatic and asymptomatic segments from leaves infected with each of the three parasites, as well as an asymptomatic segment collected from a leaf with no signs of disease. Thus, we collected a total of 252 leaf segments. All segments were of approximately equal size. The two segments collected from each symptomatic leaf were spaced at least 10 centimeters apart within the leaf. We stored each leaf segment in an individual plastic bag that was then placed on ice.

Within four hours, all segments were processed back in the laboratory. Leaf segments were washed under running DI water for 30 seconds to remove fungi that were on the surface of the leaf but not attached to the leaf. Following surface washing, leaf segments were stored in a - 80C freezer.

### DNA extraction and Sequencing

Surface-washed leaf segments were ground under liquid nitrogen with a mortar and pestle and transferred to 96-well plates for DNA extraction. DNA extraction was performed with the DNEasy PowerSoil kit according to the manufacturer’s protocol (Qiagen).

We assayed the fungal communities of tall fescue leaves by sequencing the internal transcribed spacer (ITS) region. The ITS is a region of the nuclear ribosomal RNA cistron that is often used as a DNA barcode for fungi, as it has less intraspecific variation than interspecific variation (Schoch et al., 2012). We amplified the first part of the internal transcribed spacer (ITS1) with a version of the primer set ITS1F and ITS2 modified for parallel sequencing on the Illumina MiSeq platform (White, 1990; Smith & Peay, 2014). Each 25 uL PCR reaction had 2.5 uL of 10x PCR Buffer, 3.5uL of MgCl2, 1uL of ITS1-F, 1uL of ITS2, 0.5uL of dNTPs, 0.63uL of Taq polymerase, 13.12uL of water, and 3uL of DNA. The reactions were performed with the following cycle conditions: initial denaturation 95C for 1 minute followed by 40 cycles of 94°C for 30 seconds, 52°C for 30 seconds, 68°C for 30 seconds and a final elongation at 68°C for 7 minutes. We visualized PCR products using gel electrophoresis, cleaned samples with AMPure beads (Lundberg, Yourstone, Mieczkowski, Jones, & Dangl, 2013), and concentration-normalized (using Qubit Fluorometric Quantitation, Life Technologies, Germany). The cleaned amplicons were pooled in one run on an Illumina MiSeq instrument (Illumina, San Diego, CA, USA) at the UNC High Throughput Sequencing Facility using a paired-end 2 × 250 bp kit. A spike of 10% PhiX was added to the library to increase sample heterogeneity. All raw sequence data is deposited in the National Center for Biotechnology Information Sequence Read Archive (accessions XXXXXX-XXXXXX) and data is available at (https://doi.org/10.5061/dryad.5x69p8d0v).

### Fungal Community Analysis

All statistical analyses were performed with leaf segment as the unit of observation, and in the R environment, version 3.6.0 (R Core Development Team 2012). Fungal sequences from the pooled samples were assigned to individual leaf segments (i.e. demultiplexed) using Illumina bcl2fastq pipeline (v.2.20.0), and sequencing adapters were removed from the fungal ITS sequences using cutadapt (v.1.15)(Martin, 2011). Illumina-sequenced amplicon data of microbial communities is often clustered into operational taxonomic units (OTUs) based on a fixed dissimilarity threshold. This clustering reduces the rate at which sequencing errors are misinterpreted as biological variation. The DADA2 package in R models and corrects Illumina-sequenced amplicon errors and infers exact amplicon sequence variants (herein referred to as taxa), meaning these taxa are biological variants and not sequencing noise (Callahan et al., 2016). This method can resolve biological differences at a high resolution, and the output can be directly compared between studies without the need to reprocess the pooled data. We therefore employed DADA2 in this study. Quality control of sequencing reads for each leaf segment consisted of truncating reads at the first quality score of 2 (a quality score of 2 indicates a portion of the sequence that contains mostly low-quality reads of Q15 or less), and removing any read with ambiguous base calls or greater than two expected errors. Reads shorter than 50 bases after quality trimming were removed.

### Statistical analyses: Diversity

To compare the diversity of fungal communities among asymptomatic and symptomatic leaves of tall fescue, we quantified Hill’s series of diversity. Hill’s series of diversity (Hill, 1973) is comprised of three orders (q) of diversity that summarize information about the number and relative abundances of taxa. In Hill’s series, the values of q (0, 1, 2) indicate the relative weight applied to relative abundance (Bent & Forney, 2008). We estimated fungal richness (Hills’ N0, q=0), exponentiated Shannon entropy (Hill’s N1, q=1), and inverse Simpson diversity (Hill’s N2, q=2) (Jost, 2006). Because Shannon entropy and Simpson diversity are less sensitive to the detection of rare taxa than species richness, they each place more weight on abundant taxa. Simpson diversity places even more weight on abundant taxa than Shannon diversity. The unit of each of the numbers within Hill’s series of diversity is effective number of taxa, allowing comparisons across each value of diversity. Hill’s numbers were calculated with the vegan package (version 2.5.3)(Oksanen et al. 2013).

To test whether fungal diversity is associated with symptom type, we used linear mixed models to explain Hill’s N0, N1 and N2. In order to meet assumptions of normality and homoscedasticity of the residuals, we log-transformed diversity. We included symptom type (7 levels: the asymptomatic and symptomatic segments from leaves with each of the three focal symptoms, plus asymptomatic segments from asymptomatic leaves) as fixed effects. High-throughput sequencing of pooled samples can result in samples that differ in sequencing depth; we accounted for observational bias stemming from this difference by incorporating sequencing depth into the models as another fixed effect, following Bálint et al., 2015. Leaf ID collection point were included as random effects, with leaf ID nested within collection point. Linear mixed effects models were assessed in nlme, and we used emmeans (Lenth, 2018, version 1.3.2) to evaluate the estimated marginal means of diversity indices for each explanatory variable level, adjusted for multiple comparisons (Tukey HSD). After accounting for the variation explained by random effects and sequencing depth, we compared the partial residuals of Hill’s numbers among the treatments with Tukey’s HSD.

To quantify any changes in within-host microbial diversity as disease progressed, we used disease severity (percent leaf area exhibiting symptoms) as a measure of disease progression. Specifically, for each of the three parasite symptom types, we fit three linear mixed models using R package nlme (version 3.1-142, Pinheiro, Bates, DebRoy, Sarkar, & Team, 2013) with Hill’s N1, Hill’s N2, or Hill’s N3 as the dependent variable and disease severity as an independent variable. Hill’s N1, Hill’s N2, and Hill’s N3 were log-transformed to meet assumptions of homoscedasticity and normality. Each model also included the square root of sequencing read numbers obtained for a leaf segment as an independent variable, and collection point as a random effect. Thus, each model had the following form: Hill ∼ sqrt(readNumbers) + Symptom + 1|Collection Point. A clear relationship between disease severity and fungal diversity, in the same direction as the overall association of fungal diversity and symptom type, would suggest that the parasite progressively impacts the fungal microbiome as the disease severity increases. No relationship between disease severity and fungal diversity, or a relationship in the opposite direction to the overall association of fungal diversity and symptom type, would suggest that any association between the parasite and microbiome is not due to a progressive impact of the parasite on the microbiome.

### Statistical Analyses: Community Composition

To test the hypothesis that parasites that modify niches within their host by creating necrotic tissue alter fungal community composition, we tested whether fungal community composition was correlated with symptom type. Bray-Curtis distances were calculated among leaf segments separately and visualized using non-metric multidimensional scaling (NMDS) implemented in the phyloseq package (version 1.24, Mcmurdie & Holmes, 2013). We performed a permutational MANOVA using the adonis function in the vegan package. The predictors were collection point and symptom type. The adonis function is sensitive to the order in which variables are added, so models were run multiple times, varying the order of predictors, to verify important predictors and we report predictors that were significant regardless of order (following Vannette, Leopold, & Fukami, 2016).

We also investigated whether the homogeneity of the composition of the fungal leaf microbiome varied with symptom type. For each leaf segment, we quantified the distances from each measured Bray-Curtis distance to the centroid of Bray-Curtis distance for that leaf segment’s symptom type. We then compared the dispersion of the measurements within each symptom type across categories using the betadisper function in the vegan package.

#### Statistical Analyses: Interpretation

To improve statistical inference, we report our results using the language of the “statistical clarity concept,” instead of emphasizing statistically significant results (Dushoff, Kain, & Bolker, 2019). This approach puts forward that the results of null hypothesis significance testing are most usefully interpreted as a guide to whether a certain effect can be seen clearly in a particular context.

## Results

From the 252 leaf segments, Illumina generated 6,650,600 ITS1 reads. Of these, 4,483,694 reads passed quality filtering. This represents a mean number of reads per leaf segment of 17,792. Using DADA2, we identified 2961 unique amplicon sequencing variants (taxa). This represents a mean number of taxa per leaf segment of 70.8 (median of 62). All taxa placed within the kingdom fungi. 12.5% of taxa could not be placed lower than the kingdom fungi. Of the taxa that could be placed lower than the kingdom fungi, 99.2% placed within Ascomycota (1459) or Basidiomycota (1110). At the class level, most taxa within Ascomycota were assigned to Dothideomycetes (808) or Sordariomycetes (307), and most taxa within Basidiomycota were assigned to Agaricomycetes (691) or Tremellomycetes (186). The following analyses consider these 2961 taxa delineated by DADA2.

### Diversity

After accounting for variation in sequencing depth, symptom type strongly and clearly predicted variation in all three numbers in Hill’s series (ANOVA *P* < 0.0001). There were fewer fungal taxa (Hill’s N0) and there was lower diversity (Hill’s N1 and N2) in leaf segments that exhibited symptoms of necrotrophic parasites compared to leaf segments that were asymptomatic or symptomatic of either hemibiotrophic or biotrophic parasites (Table S1, Figure 1, Tukey’s HSD, *P* < 0.01). Specifically, when comparing the mean of each of Hill’s numbers between leaf segments symptomatic of necrotrophic parasites and all other leaf segments, Hill’s N0 was 41.4-55.0% lower, Hill’s N1 was 66.3-77.6% lower, and Hill’s N2 was 58.8-71.1% lower. In contrast, there were no clear differences in fungal richness or diversity between asymptomatic leaf segments and those symptomatic of hemibiotrophic or biotrophic parasites (p>0.05). Finally, there were no clear differences in fungal richness or diversity between asymptomatic leaf segments of any type, whether from leaves with any of the parasite symptoms or leaves that were wholly asymptomatic. Thus, the leaf segments symptomatic of a necrotrophic parasite came from leaves that were not detectably different from any other leaves, which suggests that the fungal diversity of the leaves symptomatic of necrotrophic parasites was not lower than other leaves prior to infection by necrotrophic parasites.

**Figure 1.**
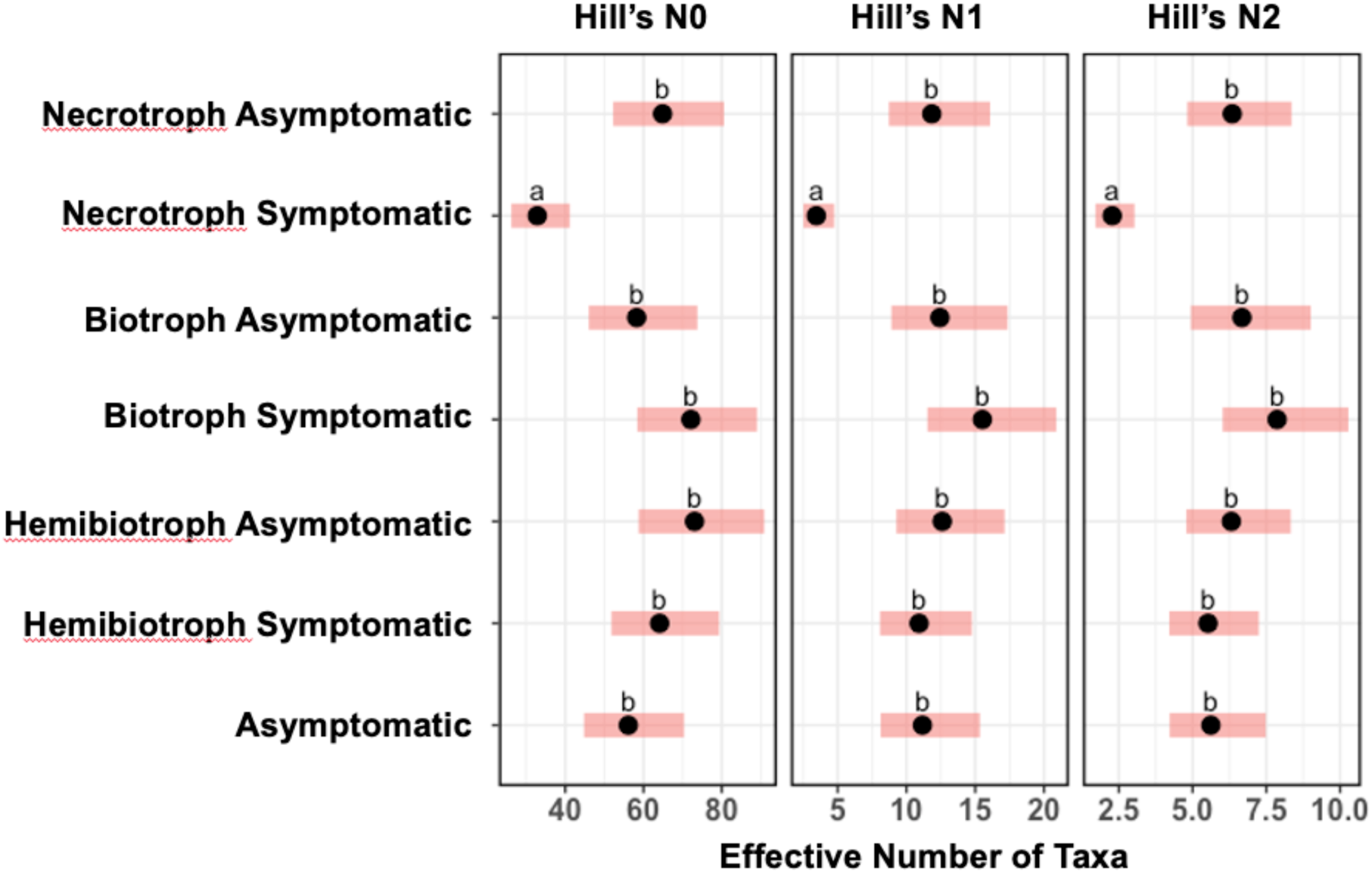
Necrotrophic symptoms were associated with foliar fungal communities that were less diverse. Panels show fungal diversity quantified using Hill numbers for observed species richness (N0), exponentiated Shannon entropy (N1), and inverse Simpson’s diversity (N2). Letters mark differences in Hill’s numbers evaluated with Tukey’s HSD. Points are estimated marginal means with red bands indicating ± 95% confidence intervals.

To investigate the lower richness and diversity of leaf segments symptomatic of necrotrophic parasites, we considered how the diversity metrics as defined in Hill’s series varied with estimated disease severity (percent leaf area exhibiting symptoms of a given parasite feeding strategy) within the segments symptomatic of necrotrophic parasites. We had predicted that if a necrotrophic parasite decreases the diversity of the fungal microbiome as it grows within its host, fungal diversity would decrease with necrotrophic parasite disease severity. Instead, fungal diversity weakly increased with necrotrophic parasite disease severity (Figure 2, Table S4; Hill’s N0, *P* = 0.010; Hill’s N1, *P* = 0.043; Hill’s N2, *P* = 0.064). Because these three correlations were not negative, these results suggest that fungal diversity, particularly richness, does not decrease progressively as a necrotrophic parasite spreads through the leaf.

**Figure 2.**
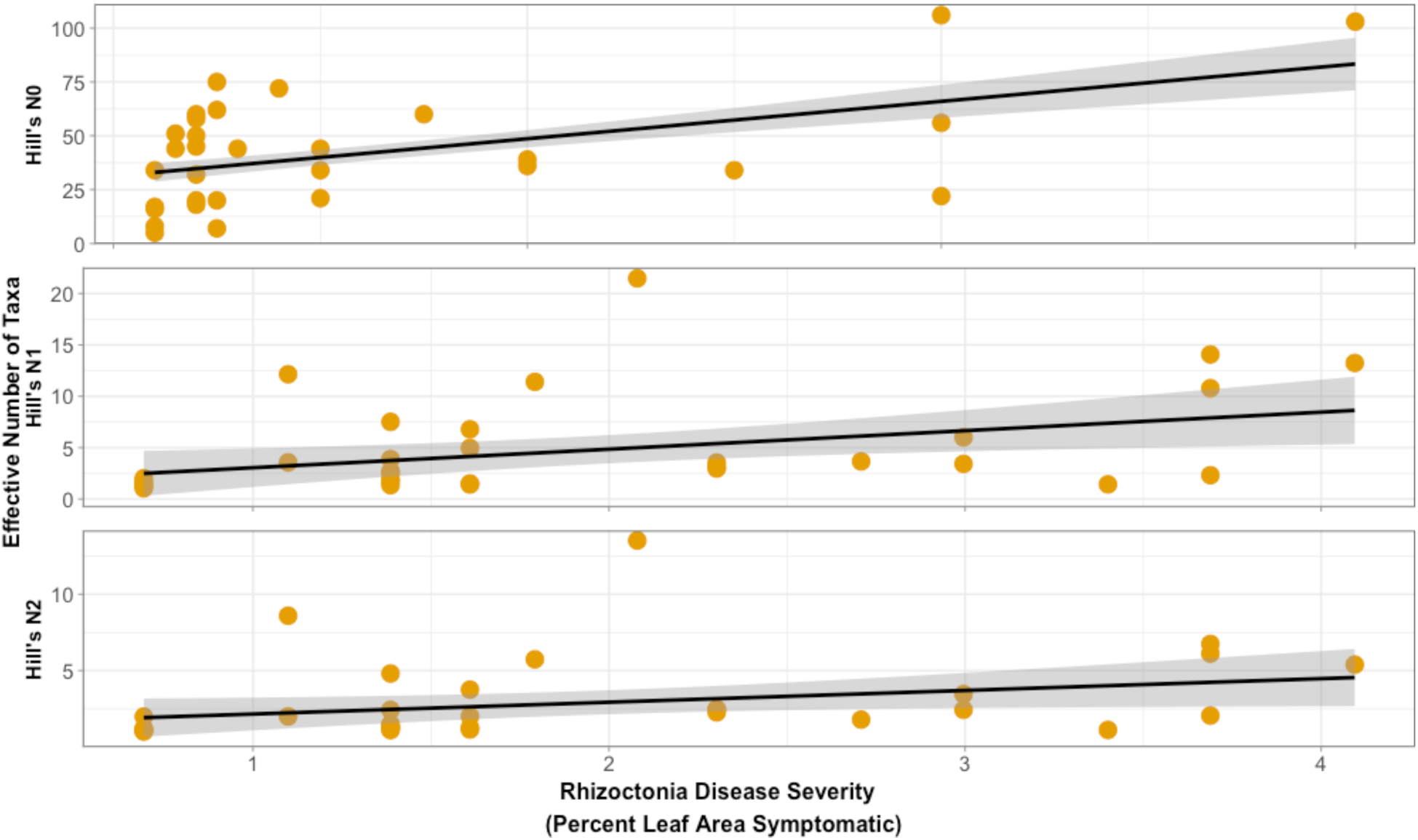
Leaf-associated fungal diversity and richness did not have a negative correlation with necrotrophic disease severity (percent leaf area exhibiting symptoms), suggesting that fungal community diversity does not change progressively as a parasite grows within a leaf. Panels show fungal diversity quantified using Hill numbers for observed species richness (N0), exponentiated Shannon entropy (N1), and inverse Simpson’s diversity (N2). Each point represents a leaf segment. Lines represent best-fit regressions between disease severity and the diversity metric.

### Community Composition

We analyzed variation in community composition using the Bray-Curtis distance metric. The fungal community composition of leaf segments with symptoms of necrotrophic parasites differed from leaf segments that were asymptomatic or symptomatic of other parasites (Figure 3, Table S2, PerMANOVA, *P* = 0.001). In other words, necrotrophic symptoms were associated with not only fewer fungal taxa, but also a different assemblage of fungal taxa compared to the other parasites. We had predicted that if a necrotrophic parasite alters the assemblage of fungal taxa as it grows within its host, fungal composition would change with estimated disease severity. Within the leaf segments that exhibited necrotrophic symptoms, disease severity did predict fungal community composition, but accounted for a small amount of variation (Figure 4, PerMANOVA, R^2^ = 0.08, *P* = 0.043). To test if any parasite symptoms were associated with a more homogeneous fungal community, we quantified beta diversity within each symptom type as the distances to the centroid from all measured Bray-Curtis dissimilarities among leaf segments of that symptom type. There was no effect of symptom type on the homogeneity of fungal community composition (Figure S9, ANOVA, *P* = 0.693). Finally, considering fine-scale variation in the fungal microbiome, within leaves symptomatic of any of the three parasite feeding strategies, the symptomatic leaf segments differed from the asymptomatic leaf segments in the relative abundance of multiple fungal genera (supplementary material).

**Figure 3.**
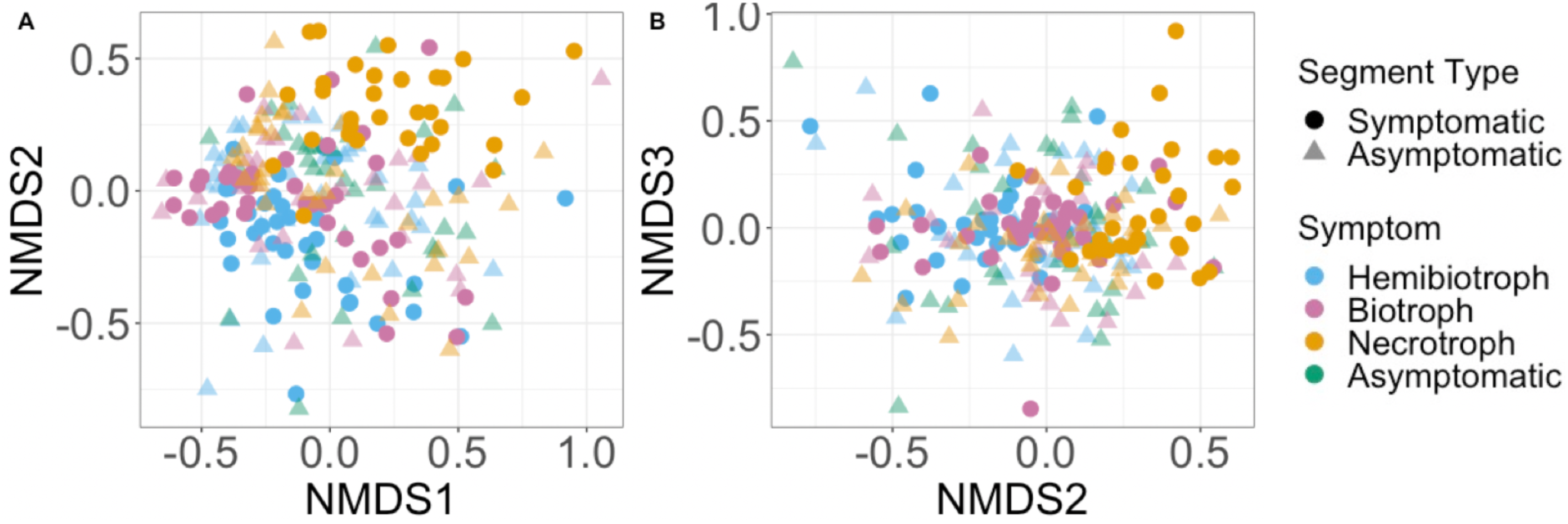
Leaf segments with necrotrophic symptoms had foliar fungal communities that differed in taxonomic composition from asymptomatic leaf segments and leaf segments with symptoms of other parasite feeding strategies. (PerMANOVA p<0.001, stress=0.18), su. Fungal taxonomic composition was quantified by the Bray-Curtis distance metric and is illustrated by non-metric multidimensional scaling (NMDS). Each point represents a leaf segment. As indicated by color, each leaf segment was either asymptomatic or symptomatic of one parasite feeding strategy (necrotroph, hemibiotroph, or biotroph).

**Figure 4.**
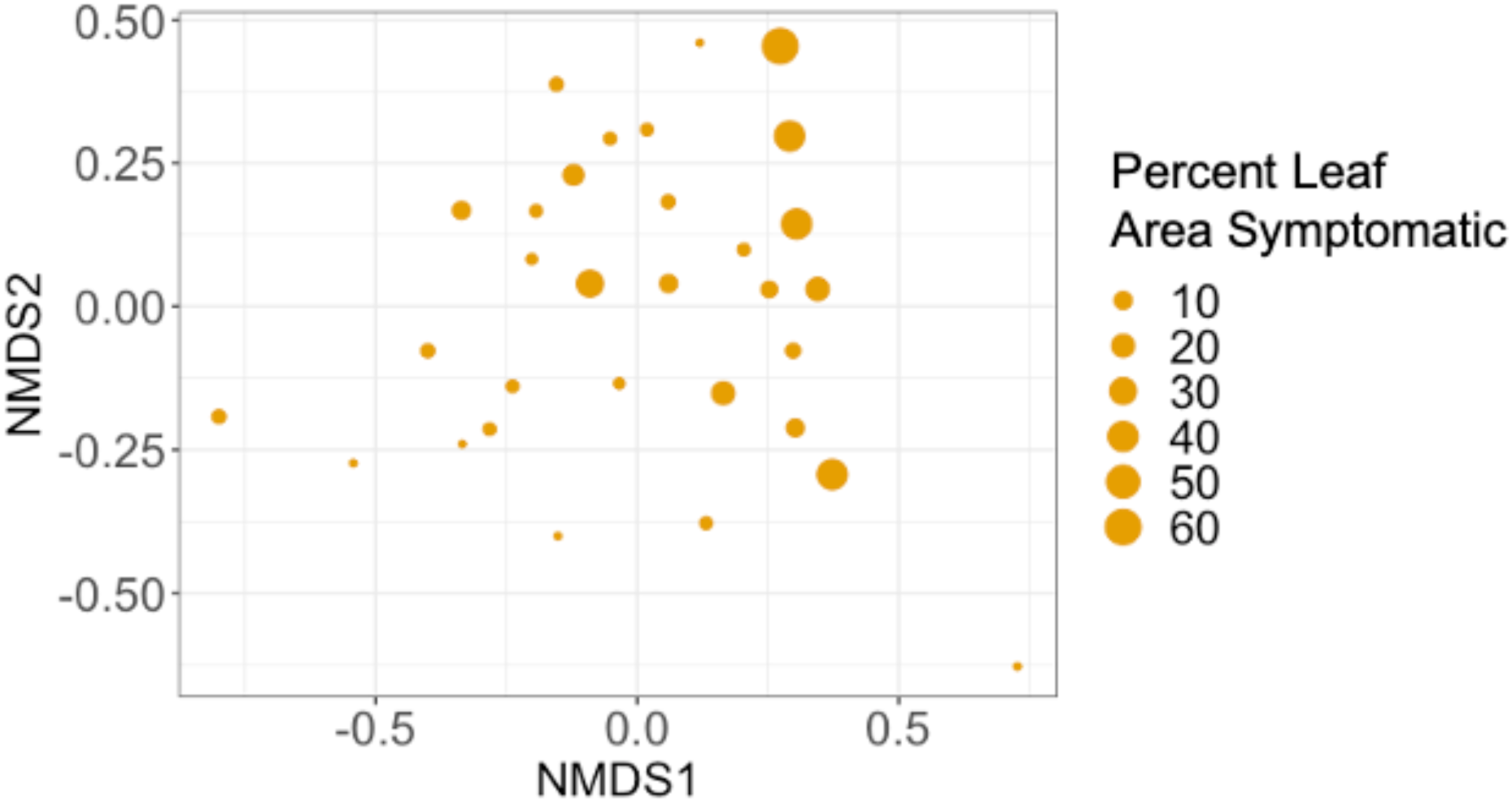
Severity of symptoms caused by necrotrophic parasites predicted fungal community composition, but only explained a modest amount of variation, suggesting that fungal community composition does not change progressively as a parasite grows within a leaf. Fungal taxonomic composition was quantified by the Bray-Curtis distance metric and is illustrated by non-metric multidimensional scaling (NMDS). Each point represents a leaf segment. Point size indicates percent leaf area symptomatic of necrotrophic parasites.

## Discussion

This study used a trait-based analysis of multiple co-occurring fungal parasites in a field population of a grass host to evaluate how parasites alter the fungal microbiome of the host. Microbiome diversity and composition were associated distinctly with symptoms of parasites with a necrotrophic feeding strategy, and not with other parasites. These results are consistent with a hypothesis based on niche modification: parasites with traits that more strongly impact the host environment and available niches within the host also more strongly impact the host microbiome.

In niche modification, a species changes the types of niches available within a site and, consequently, the identities of species that can colonize the community (Lewontin, 1983; Fukami, 2015). Among parasites infecting plant leaves, only parasites with a necrotrophic feeding strategy create necrotic host tissue throughout their entire infection process (Glazebrook, 2005; Suzuki & Sasaki, 2019). We therefore expected necrotrophs to be particularly strong niche modifiers that impact the host environment and consequently, the host fungal community. Indeed, symptoms of necrotrophic parasites were associated with fungal communities of lower diversity relative to asymptomatic leaves, while symptoms of two other types of parasite feeding strategies (biotrophs and hemibiotrophs) were not.

Our hypothesis that necrotrophic parasites are particularly strong niche modifiers was further supported by analysis of fungal community composition. The composition of the fungal communities of leaf segments exhibiting symptoms of necrotrophic parasites differed from that of asymptomatic leaves, while there was no clear difference in fungal composition between leaf segments exhibiting symptoms of biotrophic or hemibiotrophic parasites and that of asymptomatic leaves. We hypothesize that this shift in composition resulted from necrotrophic parasite infections making the host environment more suitable for saprobes (Suzuki & Sasaki, 2019).

Fungal community composition was only weakly associated with necrotrophic parasite disease severity, and fungal diversity did not have a negative correlation with disease severity. These results suggest that fungal community composition and diversity do not change progressively as a parasite grows within a leaf. The parasite may instead disrupt the host environment, and consequently, the fungal microbiome, upon initial infection. Such microbiome disruption upon initial infection is consistent with evidence from at least one other system; in experimental inoculations of frogs with *Batrachochytrium dendrobatidis*, microbiome diversity declined upon infection and had no relationship with pathogen load (Jani and Briggs, 2014).

While we expected necrotrophs to act as particularly strong niche modifiers, we expected hemibiotrophs to modify their environment as well, given that they create necrotic tissue in the latter part of the infection process (Glazebrook, 2005; Suzuki & Sasaki, 2019). However, we found contrasting results between necrotrophic and hemibiotrophic parasites; the fungal communities of leaf segments exhibiting symptoms of hemibiotrophic parasites had no clear differences in diversity and composition compared to those of asymptomatic leaf segments.

While both hemibiotrophs and necrotrophs ultimately require killing host cells, they differ in how they initially interact with host tissue. Our results therefore suggest that the initial infection by a necrotrophic parasite is the key stage in which diversity of the microbiome declines. This is consistent with a weakly supported positive correlation between disease severity and fungal taxa diversity that we observed, as diversity was lowest when necrotrophic parasite disease severity was low (i.e. change in diversity occurred early in the infection process).

While we interrogated relationships between parasites and the fungal microbiome, there is growing evidence that bacterial and fungal microbiota associate with different factors (Elhady et al., 2017; Bergelson, Mittelstrass, & Horton, 2019). For example, while we found that a biotrophic parasite had no relationship with fungal microbiome diversity, recent work investigating the bacterial microbiome of wheat found that leaves infected with a parasite infecting as a biotroph had higher bacterial diversity than uninfected leaves (Seybold et al., 2019). Such differences between the fungal and bacterial microbiome may result from their differences in generation times and abundances within a host. For more complete understanding of parasite—microbiome associations, studies that integrate surveys of the bacterial and fungal communities will be essential (Porras-alfaro & Bayman, 2011; Laforest-Lapointe & Arrieta, 2018).

In studies of plant, human, and other animal diseases, increasing numbers of studies are characterizing how microbial communities associate with specific parasites and progression of disease (Cho & Blaser, 2012; Jani & Briggs, 2014a; Jakuschkin et al., 2016; Lebreton et al., 2019). Here, we propose that functional traits can be used to explain variation among parasites in their associations with host microbiota. Trait-based approaches have played an important role in plant functional ecology (Adler et al., 2014; Cadotte, 2017). More recently, Zanne et al., 2019 advocated complementing traditional and genomic approaches to fungal functional ecology with trait-based approaches. Among traits of parasites, they discuss how parasite feeding strategy (what they refer to as nutritional strategy) can help predict how parasites interact with their abiotic and biotic environment. Integrating that trait-based prediction with the concept of niche modification, we suggest that parasite feeding strategy can determine whether the parasite is a strong niche modifier, and thus explain relationships between parasites and host-associated microbiota.

## Acknowledgements

We thank James Umbanhowar for helpful comments. We thank Anita Simha for help collecting leaves, Maggie Wagner and Posy Busby for the primers used, and Betty Wanjiru for help with the PCR amplifications. This work was supported by the NSF-USDA joint program in Ecology and Evolution of Infectious Diseases (USDA-NIFA AFRI grant 2016-67013-25762) and the University Cancer Research Fund. K.R.O. was supported by graduate research fellowships from the Triangle Center for Evolutionary Medicine and the National Science Foundation, and F.W.H. was supported by a dissertation completion fellowship from the University of North Carolina-Chapel Hill (UNC-CH) Graduate School and the W.C. Coker Fellowship in Botany from UNC-CH. Development of T-BAS 2.1 was supported by the NSF Genealogy of Life (GoLife) Program (DEB-1541418).

## Data Accessibility Statement

Upon acceptance of the manuscript, all raw sequence data will be deposited in the National Center for Biotechnology Information Sequence Read Archive, and all data sets used in this study will be submitted to Dryad.

## Author Contributions

K.R.O., F.W.H., C.D.J., I.C., and C.E.M. conceived ideas and designed methodology; K.R.O. and F.W.H. performed field collections and labwork; K.R.O. performed analyses and led writing of the manuscript. All authors contributed critically to drafts and gave final approval for publication.

## Notes

https://doi.org/10.5061/dryad.5x69p8d0v

