## Supplementary Material for "Parasites as niche modifiers for the microbiome: A field test with multiple parasites"

**Supplementary Methods**

Statistical analyses: Diversity

In the case of leaf segments symptomatic of necrotrophs, we took advantage of a five-locus phylogeny of *Ceratobasidiaceae* to investigate fungal diversity patterns excluding the taxa that may be causing the symptom (Gónzalez et al., 2016). *Ceratobasidiaceae* is a monophyletic family comprised of the genera *Ceratobasidium* and *Thanatephorus*, the teleomorphs of anamorphic *Rhizoctonia* fungi. While these fungi have been documented as saprobes and beneficial endomycorrhizal symbionts of orchids (Jiang, Lee, Cubeta, & Chen, 2015), they are parasites on grasses (Burpee & Martin, 1992). We investigated fungal diversity patterns excluding these taxa as these taxa may be the cause of observed symptoms and changes to the host-associated fungal community. Using phylogenetic placement through T-BAS (Carbone et al., 2019), we identified taxa that placed within the family (Gónzalez et al., 2016). We then created datasets in which these taxa were excluded and repeated the diversity analyses described in the main text.

**Statistical Analyses: Community Composition**

The ITS sequence for each taxon identified by DADA2 was compared against the UNITE fungal ITS database (version 7.2 release date, 28 June 2017, Kõljalg et al., 2013) via BLASTn. Additionally, sequences were evaluated for phylogenetic placement in T-BAS applying the Evolutionary Placement Algorithm (EPA) (Berger & Stamatakis, 2011) implemented in RAxML (version 8, Stamatakis, 2014) through the CIPRES RESTful application programming interface (API) (Miller et al., 2015). Briefly, query ITS sequences were first aligned to the published reference sequence alignment (TreeBASE Study S15006; Gónzalez et al., 2016) using MAFFT

(Katoh & Toh, 2010) and this comprehensive alignment was then used to map unknown taxa using the EPA in T-BAS (Carbone et al., 2017, 2019). The best placements were based on sequence matches of unknowns to reference taxa where the cumulative likelihood weights were greater than 0.96.

We assessed if the relative abundance of each taxon varied with symptom type, using the glmFit and glmLRT functions in the package edgeR following McMurdie and Holmes (2014). Specifically, for each symptom (that of necrotrophs, biotrophs, and hemibiotrophs), we performed pairwise comparisons of relative abundance of each taxon between the asymptomatic and symptomatic segments of infected leaves. For each set of analyses (one for each symptom), we filtered out low abundance taxa by excluding from analyses those taxa that occurred in fewer than 5% of leaf segments. To account for baseline variation among leaves, we used a model that included leaf ID as a predictor, as well as symptom type. To calculate normalization factors and estimate dispersion, we used the calcNormFactors, estimateGLMCommonDisp, estimateGLMTrendedDisp and estimateGLMTagwiseDisp functions, all within edgeR. We then used the output from these functions as the input for glmFit to fit a negative binomial generalized log-linear model to the abundance of each taxon. We used glmLRT to conduct taxon-wise likelihood ratio tests for a contrast between symptom types. Following McMurdie & Holmes, 2014, all tests were corrected for multiple inferences using the Benjamini-Hochberg method to control the false discovery rate (Benjamini & Hochberg, 1995). We used a false discovery rate (FDR) cutoff of 0.05.

**Supplementary Results**

<u>Diversity</u>

We identified 26 taxa that placed within the family *Ceratobasidiaceae,* which could be the cause of the necrotrophic symptom. We investigated if there were still differences in measures of fungal taxa richness and diversity between the communities of leaf segments that exhibited symptoms of necrotrophs and those of all other parasite symptom types when these taxa were excluded from analyses (as they could be the cause of such differences). After accounting for differences in sequencing depth, parasite symptom type strongly predicted variation in all three numbers in Hill's series ($p < 0.001$, Table S1, Figure S2). Across the three numbers in Hill's series, fungal diversity was lowest within leaf segments that were symptomatic of necrotrophs, though this lower fungal diversity did not have statistical support when compared to that of segments that were asymptomatic or asymptomatic and elsewhere symptomatic of biotrophs (Tukey's HSD $p > 0.05$). Specifically, when comparing the mean of each of Hill's numbers between leaf segments symptomatic of necrotrophs and leaf segments within all other parasite symptom types, Hill's N0 was 30.3-50.1% lower, Hill's N1 was 54.1-69.1% lower, and Hill's N2 was 43.7-62.0% lower. Within leaf segments symptomatic of necrotrophs, fungal diversity still had a clear positive correlation with estimated disease severity, even when excluding those taxa placed within *Ceratobasidiaceae* (Figure S3, $p < 0.05$). The association between necrotrophic symptoms and lower fungal diversity therefore holds when just considering the fungal components that are not the cause of the symptom.

Composition

Similar to diversity analyses, we repeated analyses of fungal taxa composition on a subset of taxa that excluded those that placed within *Ceratobasidiaceae*. Even after omitting *Ceratobasidiaceae* from our analysis, the taxonomic composition of leaf segments with symptoms of necrotrophs still differed from leaf segments that were asymptomatic or symptomatic of other parasites (Figure S4, PerMANOVA, p = 0.001). Furthermore, within the subset of samples that exhibited symptoms of necrotrophs, estimated disease severity was still a clear predictor of taxa composition and still only accounted for a modest amount of variation in fungal taxa composition (Figure S5, PerMANOVA, $R^2$=0.10, $P$ = 0.02). Therefore, the difference in composition was not due to the presence of taxa that were causing the symptom.

We investigated how relative abundance of the 2961 taxa identified varied between leaf segments symptomatic of each parasite and the asymptomatic segments that were infected with that parasite elsewhere. After accounting for baseline differences among leaves by including a random effect for leaf, we identified 11 taxa, representing ten genera, that clearly differed ($p < 0.05$) between leaf segments symptomatic of necrotrophs and the asymptomatic segments of leaves infected with necrotrophs (Figure S6). One taxon placed within *Ceratobasidium*, and leaf segments symptomatic of necrotrophs had higher relative abundance of this taxon than the asymptomatic segments (log fold change of 9.40 (variant 153). We identified 14 taxa, representing 11 genera, that clearly differed between leaf segments symptomatic of biotrophs and the asymptomatic segments of leaves infected with biotrophs (Figure S7). In contrast, we identified 45 taxa, representing 35 genera, that differed between leaf segments symptomatic of hemibiotrophs and the asymptomatic segments of leaves infected with hemibiotrophs (Figure S8).

**Table S1: Variation of Hill's series of diversity, explained by linear mixed models.** Hill's N0 is species richness; N1 is the antilogarithm of Shannon's diversity, and N2 is the inverse of Simpson's Diversity

| | Hill's N0 | | | | Hill's N1 | | | | Hill's N2 | | | |
|---|---|---|---|---|---|---|---|---|---|---|---|---|
| | F | denDF | numDF | P | F | denDF | numDF | P | F | denDF | numDF | P |
| *Fixed Effects* | | | | | | | | | | | | |
| Read numbers | 69.626 | 84 | 1 | <.0001 | 6.3416 | 84 | 1 | 0.0044 | 5.032 | 84 | 1 | 0.0275 |
| Symptom Type | 6.908 | 84 | 6 | <.0001 | 10.9944 | 84 | 6 | <.0001 | 9.317 | 84 | 6 | <.0001 |

**Table S2: Leaf-associated fungal diversity and richness had a positive correlation with estimated *Rhizoctonia* disease severity (percent leaf damaged).**

| | Hill's N0 | | | | Hill's N1 | | | | Hill's N2 | | | |
|---|---|---|---|---|---|---|---|---|---|---|---|---|
| | F | denDF | numDF | P | F | denDF | numDF | P | F | denDF | numDF | P |
| *Fixed Effects* | | | | | | | | | | | | |
| Read numbers | 9.87214 | 35 | 1 | 0.0039 | 1.97292 | 84 | 1 | 0.1711 | 1.39015 | 35 | 1 | 0.2483 |
| Nectrophic Severity | 7.64181 | 35 | 1 | 0.01 | 4.50003 | 84 | 1 | 0.0429 | 3.72272 | 35 | 1 | 0.0639 |

**Table S3: Variation of Hill's series of diversity, explained by linear mixed models, for the subset of variants that do not place within Ceratobasidiaceae.** Hill's N0 is species richness; N1 is the antilogarithm of Shannon's diversity, and N2 is the inverse of Simpson's Diversity

| | Hill's N0 | | | | Hill's N1 | | | | Hill's N2 | | | |
|---|---|---|---|---|---|---|---|---|---|---|---|---|
| | **F** | **denDF** | **numDF** | **P** | **F** | **denDF** | **numDF** | **P** | **F** | **denDF** | **numDF** | **P** |
| *Fixed Effects* | | | | | | | | | | | | |
| Read numbers | 87.686 | 84 | 1 | <.0001 | 5.9168 | 84 | 1 | 0.0171 | 1.735 | 84 | 1 | 0.1914 |
| Symptom Type | 6.259 | 84 | 6 | 0.002 | 5.6337 | 84 | 6 | 0.001 | 4.8104 | 84 | 6 | 0.0003 |

**Table S4: Variation in the composition of the fungal microbiome explained by parasite and symptom.** Results of adonis (permutational MANOVA). Predictors were included in the order indicated below. However, relative importance remained unchanged among different permutations. Values indicate the $R^2$

| | **denDF** | **numDF** | **$R^2$** | **p** |
|---|---|---|---|---|
| *Fixed Effects* | | | | |
| Read numbers | 251 | 1 | 0.008 | 0.011 |
| Symptom Type | 246 | 6 | 0.101 | 0.001 |

**Figure S1. Map of the transects in the study.** Mapped area is a grass-dominated field that is part of the Duke Forest Teaching and Research laboratory in Orange County, North Carolina. Green lines indicate the approximate location of each transect across the field, and each transect was at least 20 meters apart. The inset indicates how each transect was divided into sampling sites. Samples were collected every 20 meters along a 100 meter transect.

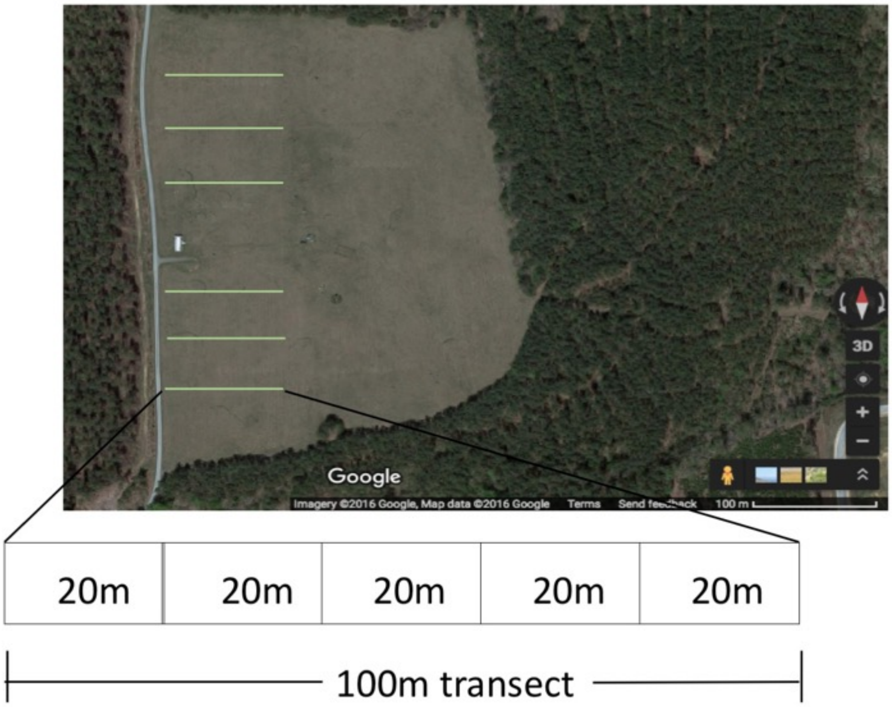

**Figure S2 When excluding taxa that did not place within *Ceratobasidiaceae*, necrotrophic symptoms were still associated with foliar fungal communities that were less diverse** . Panels show fungal diversity quantified using Hill numbers for observed species richness (N0), exponentiated Shannon entropy (N1), and inverse Simpson's diversity (N2). Letters mark differences in Hill's numbers evaluated with Tukey's HSD. Points are estimated marginal means with red bands indicating ± 95% confidence intervals.

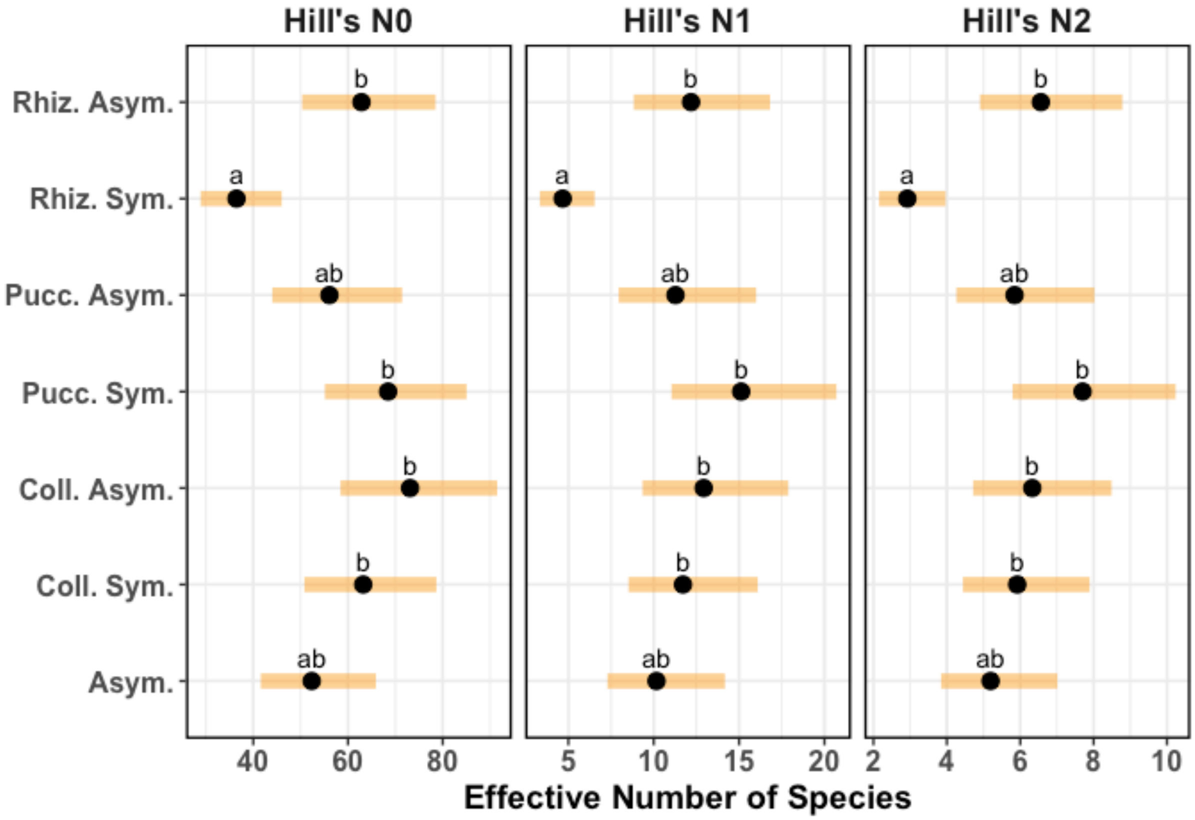

**Figure S3. After excluding taxa placed within *Ceratobasidiaceae*, Leaf-associated fungal diversity and richness did not have a negative correlation with necrotrophic disease severity (percent leaf area exhibiting symptoms), suggesting that fungal community diversity does not change progressively as a parasite grows within a leaf.**
Panels show fungal diversity quantified using Hill numbers for observed species richness (N0), exponentiated Shannon entropy (N1), and inverse Simpson's diversity (N2). Each point represents a leaf segment. Lines represent best-fit regressions between disease severity and the diversity metric.

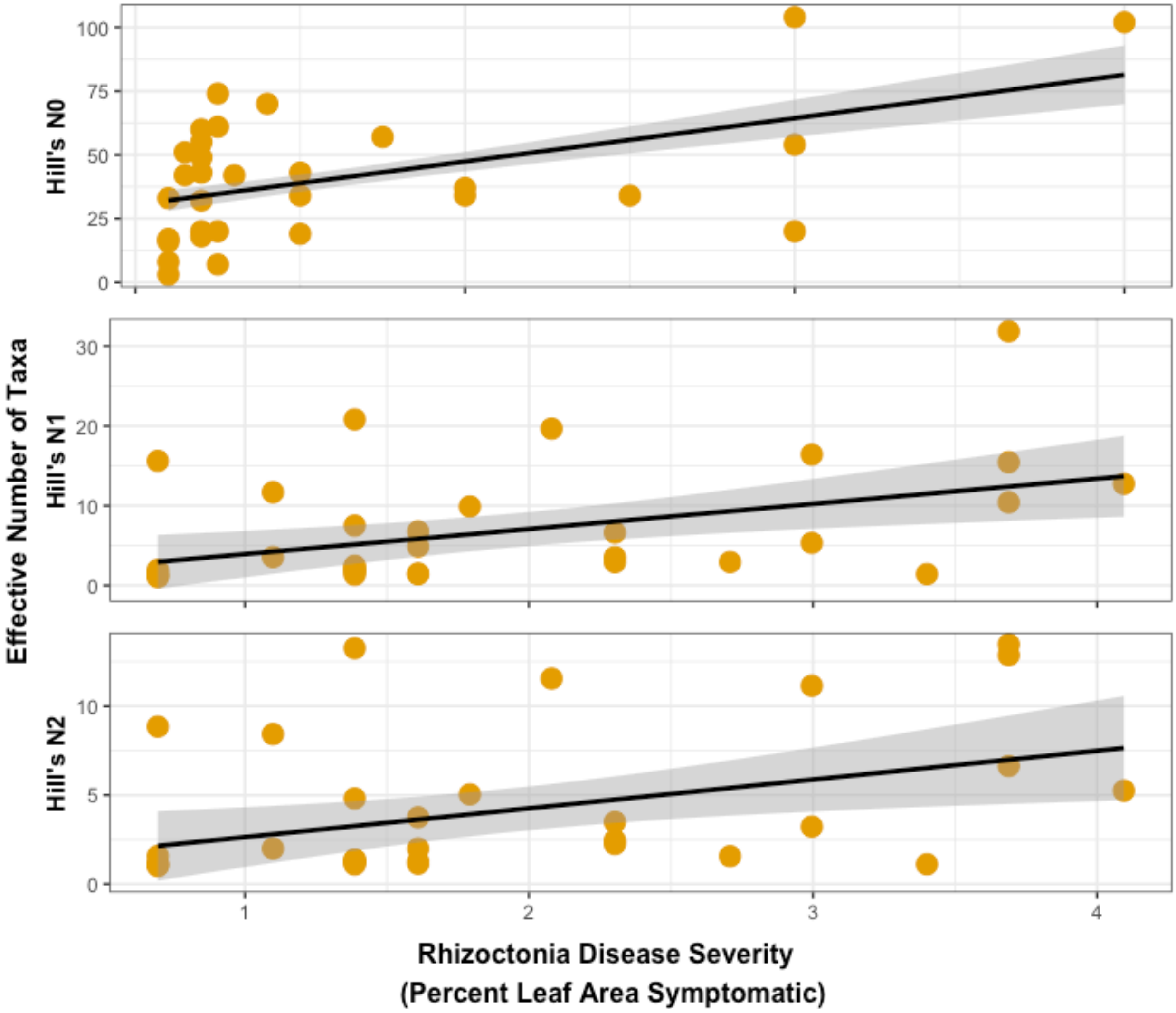

**Figure S4. Excluding those taxa that placed within *Ceratobasidiaceae*, leaf segments with necrotrophic symptoms had foliar fungal communities that differed in taxonomic composition from asymptomatic leaf segments and leaf segments with symptoms of other parasite feeding strategies,** (PerMANOVA $p<0.001$, stress=0.18)**.** Fungal taxonomic composition was quantified by the Bray-Curtis distance metric and is illustrated by non-metric multidimensional scaling (NMDS). Each point represents a leaf segment. As indicated by color, each leaf segment was either asymptomatic or symptomatic of one parasite feeding strategy (necrotroph, hemibiotroph, or biotroph).

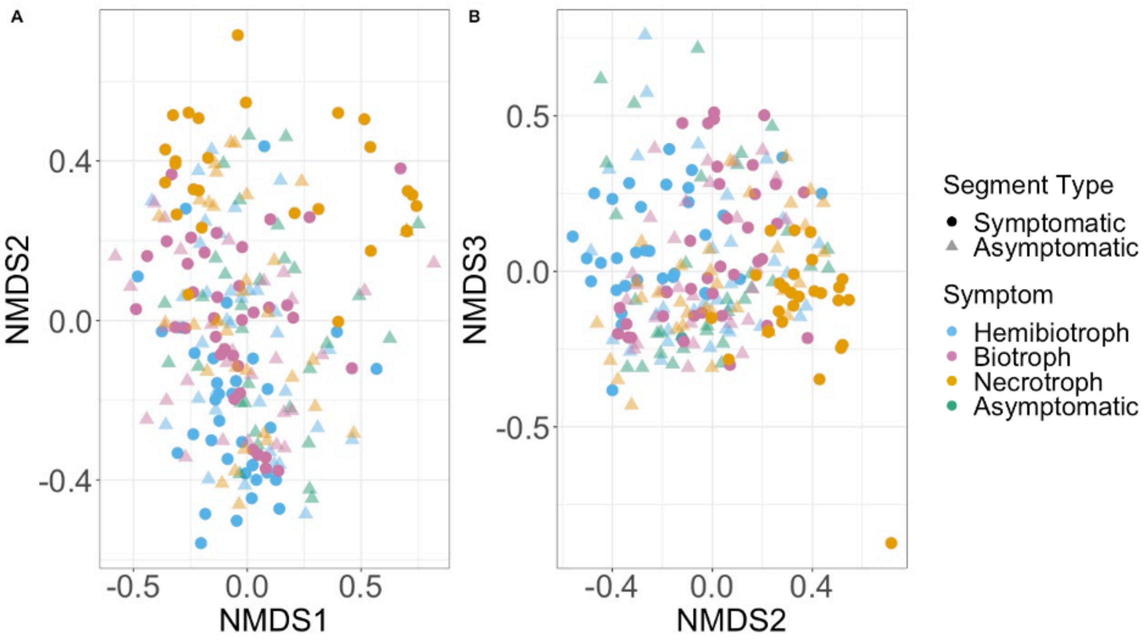

**Figure S5. Excluding taxa that placed within *Ceratobasidiaceae*, severity of symptoms caused by necrotrophic parasites predicted fungal community composition, but only explained a modest amount of variation, suggesting that fungal community composition does not change progressively as a parasite grows within a leaf.** Fungal taxonomic composition was quantified by the Bray-Curtis distance metric and is illustrated by non-metric multidimensional scaling (NMDS). Each point represents a leaf segment. Point size indicates percent leaf area symptomatic of necrotrophic parasites.

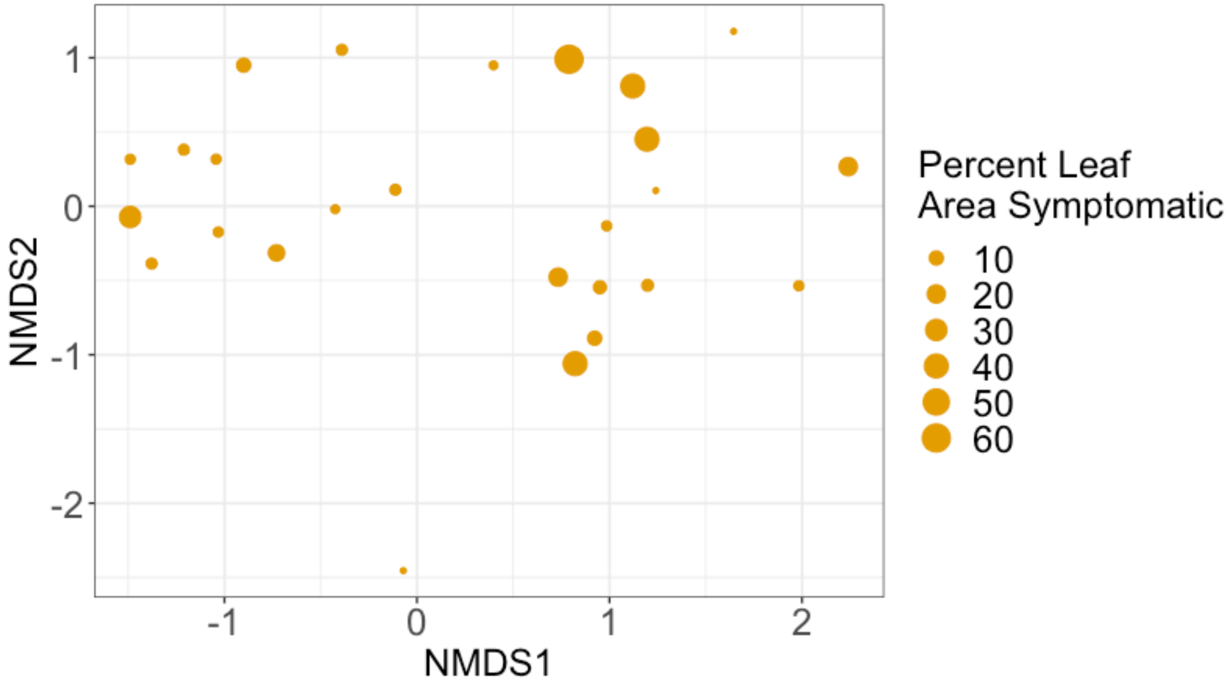

**Figure S6: 11 taxa, representing 10 genera, differed in relative abundance between the asymptomatic and symptomatic segments of leaves exhibiting symptoms of necrotrophic parasites.** Results of edgeR analysis comparing relative sequence abundance for fungal sequencing taxa between the asymptomatic and symptomatic segments of leaves exhibiting symptoms of biotrophic parasites. Bars represent the log fold difference in relative sequence abundance for taxa that differed between asymptomatic and symptomatic segments ($p<0.05$) with the genus name indicated on the x-axis.

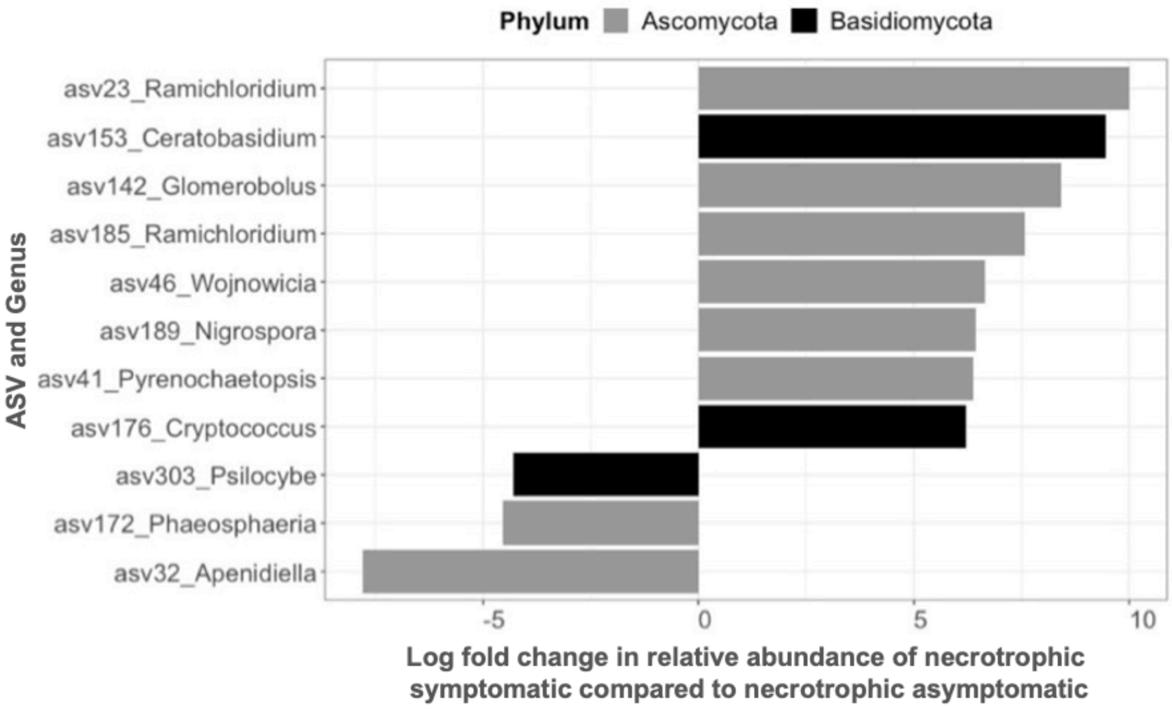

**Figure S7: 14 taxa, representing 11 genera, differed significantly in relative abundance between the asymptomatic and symptomatic segments of leaves exhibiting symptoms of biotrophic parasites.** Results of edgeR analysis comparing relative sequence abundance for fungal sequencing taxa between the asymptomatic and symptomatic segments of leaves exhibiting symptoms of biotrophic parasites. Bars represent the log fold difference in relative sequence abundance for taxa that differed significantly between asymptomatic and symptomatic segments ($p<0.05$) with the genus name indicated on the x-axis after filtering out variants that occurred in less than 5% of samples.

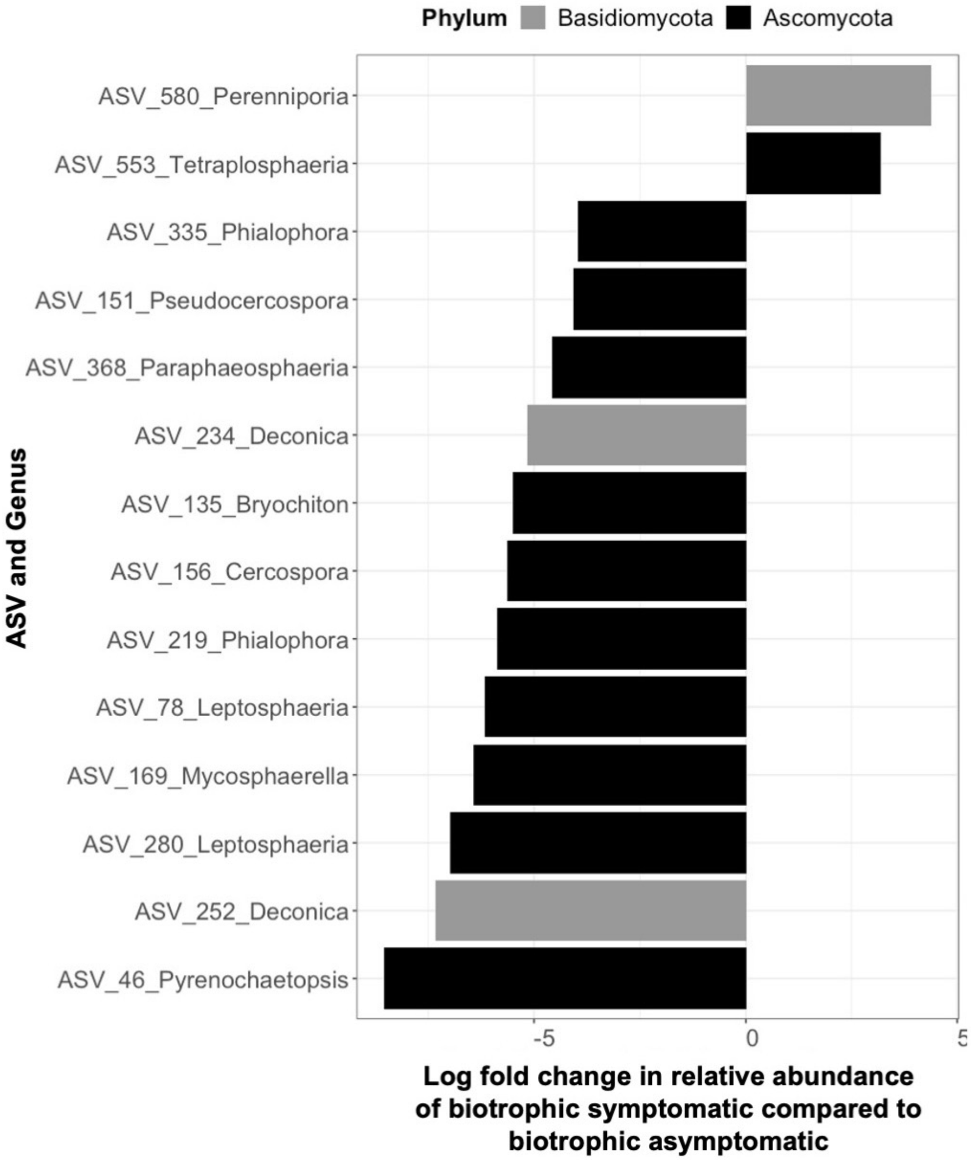

**Figure S8: 45 taxa, representing 35 genera, differed significantly in relative abundance between the asymptomatic and symptomatic segments of leaves exhibiting symptoms of hemibiotrophic parasites.** Results of edgeR analysis comparing relative sequence abundance for fungal sequencing taxa between the asymptomatic and symptomatic segments of leaves exhibiting symptoms of hemibiotrophic. Bars represent the log fold difference in relative sequence abundance for taxa that differed significantly between asymptomatic and symptomatic segments ($p<0.05$) with the genus name indicated on the x-axis after filtering out variants that occurred in less than 5% of samples.

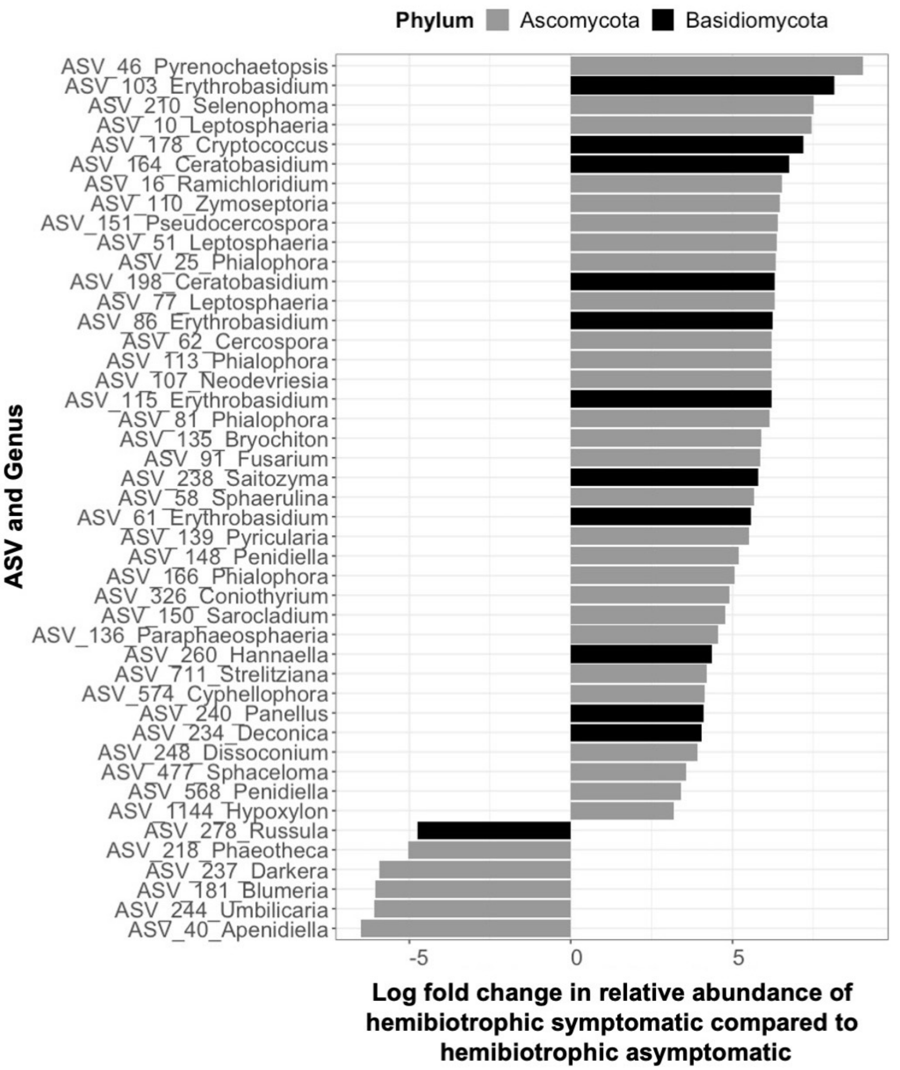

**Figure S9. There is no clear effect of parasite symptom category on the homogeneity of the composition of the fungal microbiome.** Points are measurements of the distances from measured Bray-Curtis dissimilarity to centroid of Bray-Curtis dissimilarity for a given parasite symptom category. Boxplots show the median and the lower and upper hinges correspond to the first and third quartiles. The whiskers extend from lower and upper hinges to the smallest and largest values, respectively, no further than 1.5 times the inter-quartile range from the respective hinges, respectively. All points beyond the whiskers are outliers.

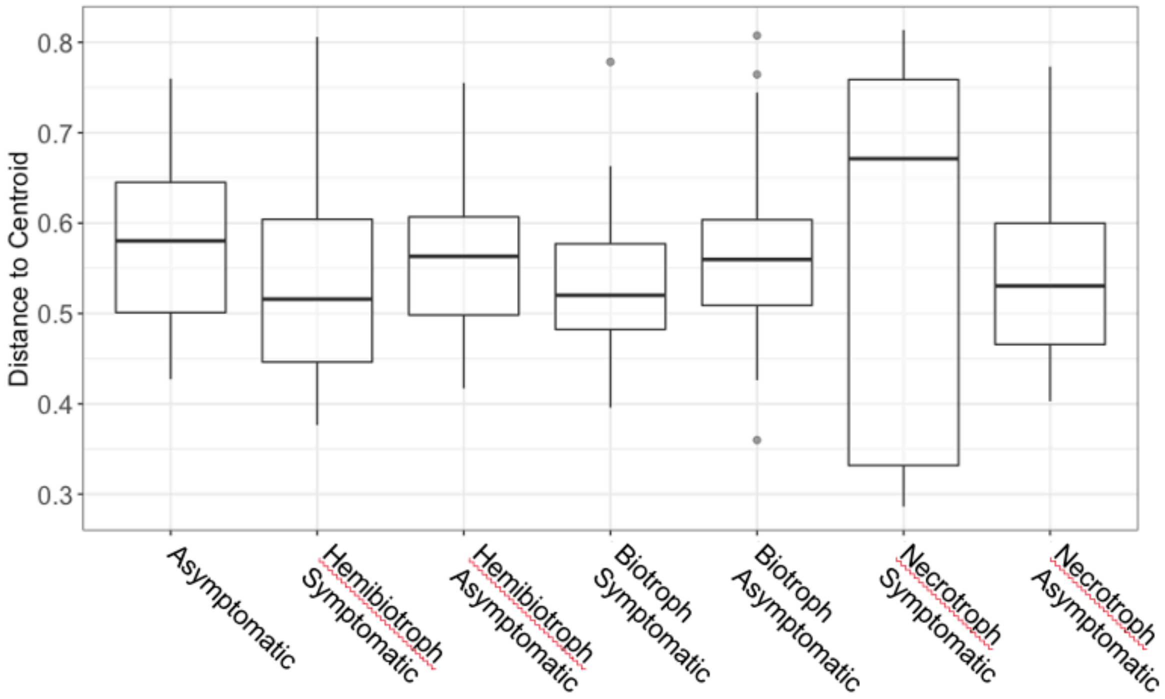

Berger, S. A., & Stamatakis, A. (2011). Aligning short reads to reference alignments and trees. *Bioinformatics*, *27*(15), 2068–2075. doi:10.1093/bioinformatics/btr320

Burpee, L., & Martln, B. (1992). Biology of Rhizoctonia Species Associated with Turfgrasses. *Plant Disease*, *76*(2), 112–117.

Carbone, I., White, J. B., Miadlikowska, J., Arnold, A. E., Miller, M. A., Kauff, F., … May, G. (2017). T-BAS : Tree-Based Alignment Selector toolkit for phylogenetic-based placement , alignment downloads and metadata visualization : an example with the Pezizomycotina tree of life, *33*(January 2018), 1160–1168. doi:10.1093/bioinformatics/btw808

Carbone, I., White, J. B., Miadlikowska, J., Arnold, A. E., Miller, M. A., Magain, N., … Lutzoni, F. (2019). T-BAS Version 2.1: Tree-Based Alignment Selector Toolkit for Evolutionary Placement of DNA Sequences and Viewing Alignments and Specimen Metadata on Curated and Custom Trees. *Microbiology Resource Announcements*, *8*(29), e00328-19. doi:10.1128/MRA.00328-19

Gónzalez, D., Rodriguez-Carres, M., Boekhout, T., Stalpers, J., Kuramae, E. E., Nakatani, A. K., Vilgalys, R., Cubeta, M. A. (2016). Phylogenetic relationships of Rhizoctonia fungi within the Cantharellales. *Fungal Biology*, *120*(4), 603–619. doi:https://doi.org/10.1016/j.funbio.2016.01.012

Jiang, J.-H., Lee, Y.-I., Cubeta, M. A., & Chen, L.-C. (2015). Characterization and colonization of endomycorrhizal Rhizoctonia fungi in the medicinal herb Anoectochilus formosanus (Orchidaceae). *Mycorrhiza*, *25*(6), 431–445. doi:10.1007/s00572-014-0616-1

Katoh, K., & Toh, H. (2010). Parallelization of the MAFFT multiple sequence alignment program. *Bioinformatics (Oxford, England)*, *26*(15), 1899–1900. doi:10.1093/bioinformatics/btq224

Kõljalg, U. *et al*. Towards a unified paradigm for sequence-based identification of fungi. *Mol. Ecol*. **22**, 5271–5277 (2013).

Miller, M. A., Schwartz, T., Pickett, B. E., He, S., Klem, E. B., Scheuermann, R. H., … O'Leary, M. A. (2015). A RESTful API for Access to Phylogenetic Tools via the CIPRES Science Gateway. *Evolutionary Bioinformatics Online*, *11*, 43–48. doi:10.4137/EBO.S21501

Stamatakis, A. (2014). RAxML version 8: a tool for phylogenetic analysis and post-analysis of large phylogenies. *Bioinformatics (Oxford, England)*, *30*(9), 1312–1313. doi:10.1093/bioinformatics/btu033
